# Structural and functional basis of proton-independent transition metal import by a canonical bacterial Nramp transporter

**DOI:** 10.64898/2026.01.27.701978

**Authors:** Shamayeeta Ray, Samuel P. Berry, Rachelle Gaudet

**Author notes:** Present address: IOCB Boston, Cambridge, MA 02139 USA.

## Abstract

Natural resistance-associated macrophage proteins (Nramps) are divalent transition metal transporters found in most organisms, typically coupling metal uptake to proton co-transport. How this coupling evolved, however, remains unclear. We present structural, functional, and evolutionary analyses of a clade B Nramp from the gut bacterium *Bacteroides fragilis* (BfraNramp). Phylogenetic reconstruction positions clade B as the most basal group of canonical Nramps, retaining conserved metal-binding motifs while lacking most residues that form the canonical proton pathway. We show that BfraNramp efficiently transports Mn²⁺ and Cd²⁺ with high apparent affinity but without proton co-transport or dependence on membrane potential or pH. Structures of metal-free and Mn²⁺-bound BfraNramp reveal an inward-open conformation and a distinct metal coordination geometry involving a conserved glutamate on transmembrane helix 3. Together, these results identify clade B Nramps as proton-independent transition metal uniporters and suggest that proton coupling emerged later in Nramp evolution, following establishment of the metal-binding site.

## Introduction

Natural resistance-associated macrophage proteins (Nramps) are a family of highly conserved membrane-bound transporters that mediate the import of essential redox-active divalent transition metals, Fe^2+^ and Mn^2+^, into cells, thus ensuring their homeostasis^1,2^. Nramps are ubiquitously present in organisms from higher eukaryotes to prokaryotes^1,3,4^. For all biochemically characterized Nramps, efficient metal transport across the cell membrane is facilitated by co-transport of protons^2,5–8^.

Based on evolutionary analysis and recent structures of multiple Nramps across bacteria and humans, Nramps share a common LeuT fold, comprising ten core transmembrane helices that are pseudo-symmetrically related^1,8–10^. In LeuT-fold transporters, transmembrane helices (TMs) 1, 2, 6, and 7 form a bundle that generally rotates relative to a rigid hash (TMs 3, 4, 8, and 9), supported by flexible arms (TMs 5 and 10)^11^. Multiple structures in different conformations reveal that the transition metal ion substrate is bound between the unwound regions of TMs 1 and 6^8–10,12,13^. Proton co-transport occurs via a pathway formed by charged residues in TMs 3 and 9^2,5,13,14^. The overall metal transport cycle of Nramps relies on the canonical alternating access between outward-open and inward-open conformations, mediated through an intermediate occluded state in which the metal ion substrate is inaccessible to solvent from both sides of the membrane^2,12,13^. This global mechanism and the associated energetics of metal and proton transport by various Nramps is similar in prokaryotes and eukaryotes^2,10,12^.

The prokaryotic Nramps have been further subdivided based on sequence similarity into three major clades, A, B and C^1,3,4^. Structural and functional studies on clade A and clade C Nramps reveal a similar proton-assisted metal transport mechanism, with conserved sequence motifs for both proton and metal binding and transport that are also present in most eukaryotes^5,8,10,12,13^. We annotate the bacterial clades A, B, C and the eukaryotic Nramps as the ‘canonical’ Nramps. Sequence-guided evolutionary studies have identified a highly diverse group of Nramp-related proteins lacking the conserved metal-binding and proton-transport residues of canonical Nramps^3,4,15^. The Nramp-related protein EleNRMT has an overall structure similar to canonical Nramps but can transport Mg^2+^, an alkaline earth metal not transported by canonical Nramps^15^. In contrast, very little is known about the clade B Nramps, the smallest clade of canonical prokaryotic Nramps and the least conserved in sequence^3,4^. Clade B Nramps are mostly present in anaerobic bacteria, including human gut microbes^3,4^. Thus, studying the clade B Nramps is important to understand how gut microbes maintain cellular transition metal homeostasis in an oxygen-depleted environment.

Here we present the structure of a clade B Nramp from the common anaerobic gut bacterium, *Bacteroides fragilis*, which we refer to as BfraNramp, in an inward-open conformation. Using detailed sequence-guided evolutionary analysis along with proteoliposome-based metal and proton transport assays, we find that clade B acts as an intermediate clade sharing functional similarity and dissimilarity with both canonical Nramps and Nramp-related proteins with respect to metal and proton transport behavior. Like the well-characterized clade A member *Deinococcus radiodurans*, DraNramp^9,11–14,16^, BfraNramp transports both Mn^2+^ and Cd^2+^ with similar efficiency. However, unlike DraNramp and other canonical Nramps^2^, BfraNramp is only minimally sensitive to pH and membrane potential and does not co-transport protons. We also resolve a Mn^2+^-bound structure of BfraNramp in which the metal ion resides between TMs 1 and 6 but with a coordination sphere distinct from other canonical Nramps, and we corroborate the importance of the Mn^2+^-interacting residues for transport using mutagenesis. Overall, our results indicate that clade B likely represents the ancestral group of transition metal ion-selective Nramps and that proton co-transport likely arose later with the introduction of polar and charged residues on TMs 3 and 9 of the LeuT-fold hash.

## Results

### The Nramp proton pathway evolved after the conserved transition metal specificity motif

To understand the evolutionary position of clade B relative to other Nramp clades, we built a rooted phylogeny of Nramps and Nramp-related proteins. We first generated a sequence alignment from a superimposed set of known and predicted Nramp structures to three members of the structurally homologous neurotransmitter-sodium symporter (NSS) transporters, which are suggested as the most closely related members of the Amino Acid-Polyamine-Organocation (APC) superfamily to the Nramps^17^. We used this structure-based sequence alignment as a seed to collect a highly diverse alignment of 97,939 sequences and build phylogenies. To obtain confident posterior probability estimates of tree topologies, we subsampled the tree to 69 sequences and sampled a posterior distribution of 100 trees across the nodes. In every sampled tree, the Nramp-related proteins formed a paraphyletic group with respect to the canonical Nramps, and clade B is basal to these canonical Nramps (Fig. 1a and Supplementary Fig. 1a). To generate sequence motifs for each clade, we then built a larger maximum-likelihood tree based on 36,700 Uniref90-filtered sequences (Supplementary Fig. 1b). While Nramp-related proteins have neither the known conserved substrate-specificity motif on TMs 1 and 6 nor the proton-pathway residues on TMs 3 and 9, clade B Nramps uniquely have all the conserved substrate-specificity motifs but are missing most of the proton-pathway residues (Fig. 1b-c). By parsimony, this suggests that the conserved metal-binding site evolved before the proton pathway except for the conserved glutamate at position 134 in DraNramp, which is also conserved as a glutamate or aspartate in many Nramp-related proteins. Of note, the TM1 aspartate and a TM6 histidine (D56 and H232 in DraNramp, respectively), are implicated both in forming the metal substrate binding site and in proton flux and are conserved in clade B^5,13,14^. This raises several questions, including: Are the clade B Nramp metal binding and transport mechanisms the same as in other canonical Nramps? And do clade B Nramps transport protons like other canonical Nramps, perhaps using an alternate mechanism, as previously hypothesized^4^?

**Figure 1.**
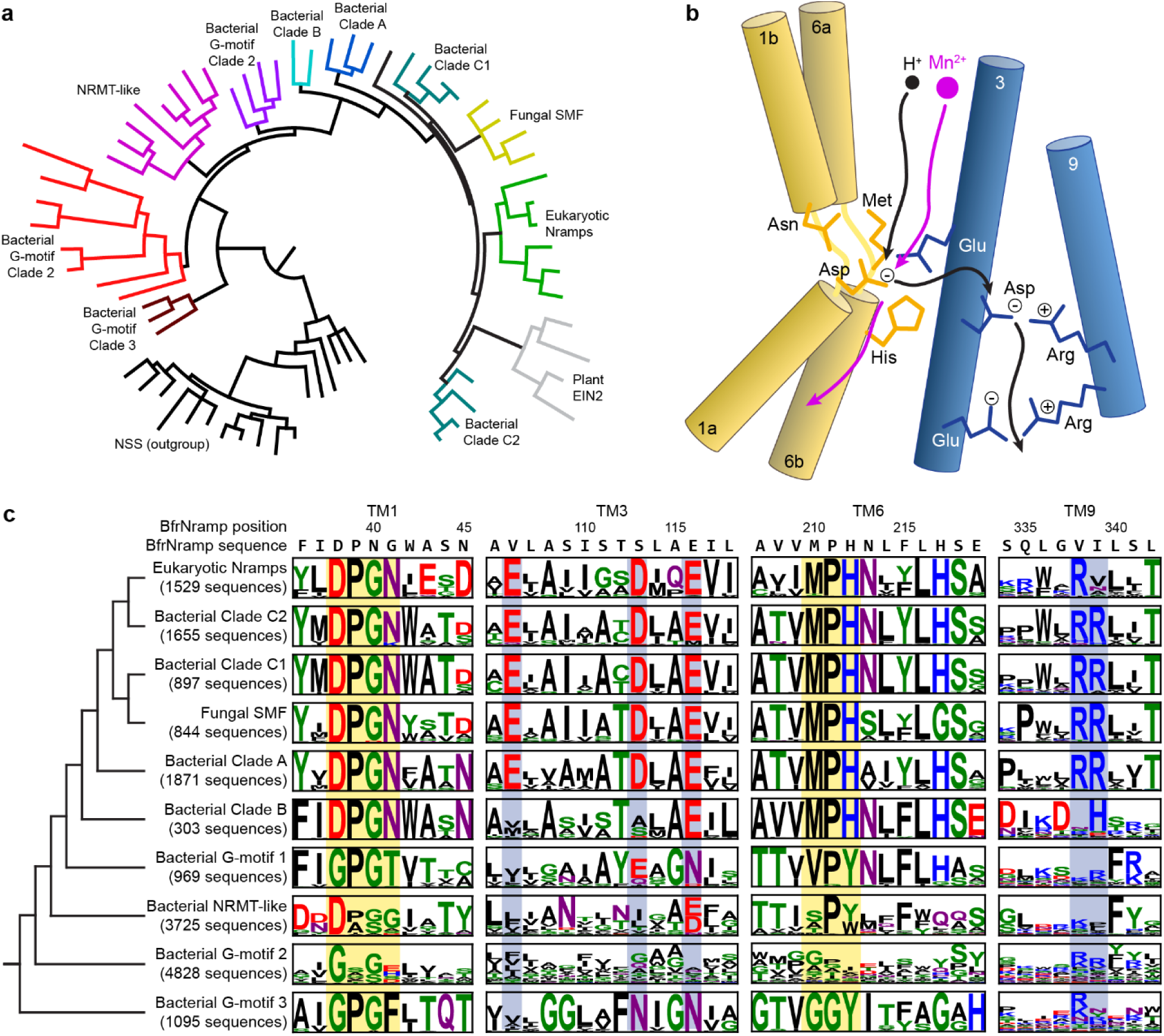
Evolution of the conserved metal-binding motif and proton pathway in Nramp homologs. (**a**) The most probable tree of 69 representative Nramp, Nramp-related and NSS transporters, with different groups highlighted. The “NRMT-like” group contains the characterized Nramp-related magnesium transporter EleNRMT^15^ and the “G-motif 3” group contains the hypothesized *B. subtilis* Mn^2+^ importer MntG^38^; the other Nramp-related groups are entirely uncharacterized. (**b**) Cartoon of a canonical Nramp, showing the key conserved metal-binding residues on bundle TMs 1 and 6 (yellow) and the conserved proton-pathway residues on hash TMs 3 and 9 (blue). (**c**) Sequence motifs of clades from a full tree and alignment of 36,700 Nramps highlighting TMs 1, 3, 6 and 9. Residues are colored according to chemical properties and heights represent information in bits. Positions illustrated in panel b are highlighted in yellow and blue.

### BfraNramp, a clade B Nramp, transports metals independent of external voltage and pH

To assess the metal transport behavior of clade B Nramps, we chose BfraNramp from the anaerobic gut bacterium *Bacteroides fragilis* (Supplementary Fig. 2a). We reconstituted BfraNramp into proteoliposomes^12,15^ and tested transport of two metal substrates: Mn^2+^, a key substrate of canonical bacterial Nramps; and Cd^2+^, a toxic metal known to bind Nramps at the substrate site^8,12^ and be transported similarly as Mn^2+^ *in vitro*^12–14^. At –120 mV and pH 7.0 (approximating physiological conditions in bacterial cell environments), BfraNramp transports both Mn^2+^ and Cd^2+^ in a concentration-dependent manner (Supplementary Fig. 2b-c). BfraNramp transports both metals with higher apparent affinity than DraNramp (∼23 times lower K_M_ for Mn^2+^, and ∼80 times lower for Cd^2+^; Table 1 and Fig. 2a-b). Further, BfraNramp transport activity was not affected by the membrane potential (0 through –120 mV) for both Mn^2+^ (Fig. 2c) and Cd^2+^ (Supplementary Fig. 2d). In contrast and as expected^13,14^, DraNramp transport activity is highly (Mn^2+^) or somewhat (Cd^2+^) enhanced as the membrane potential decreases (Fig. 2d and Supplementary Fig. 2e). Because *B. fragilis* primarily resides in the somewhat acidic colon of human gut (pH 5.5-6.5)^18^, we tested whether a change in extraliposomal pH from 7.0 to 5.5 alters the metal transport rate of BfraNramp. BfraNramp’s Mn^2+^ and Cd^2+^ transport rates showed no pH dependence (Fig. 2e and Supplementary Fig. 2f) unlike DraNramp, which as expected^14^ showed an increased Mn^2+^ transport rate at more acidic pH (Fig. 2f) but a pH-independent Cd^2+^ transport rate (Supplementary Fig. 2g). These findings indicate that BfraNramp is a robust divalent transition metal ion transporter with rates unperturbed by a range of external environmental conditions.

**Figure 2.**
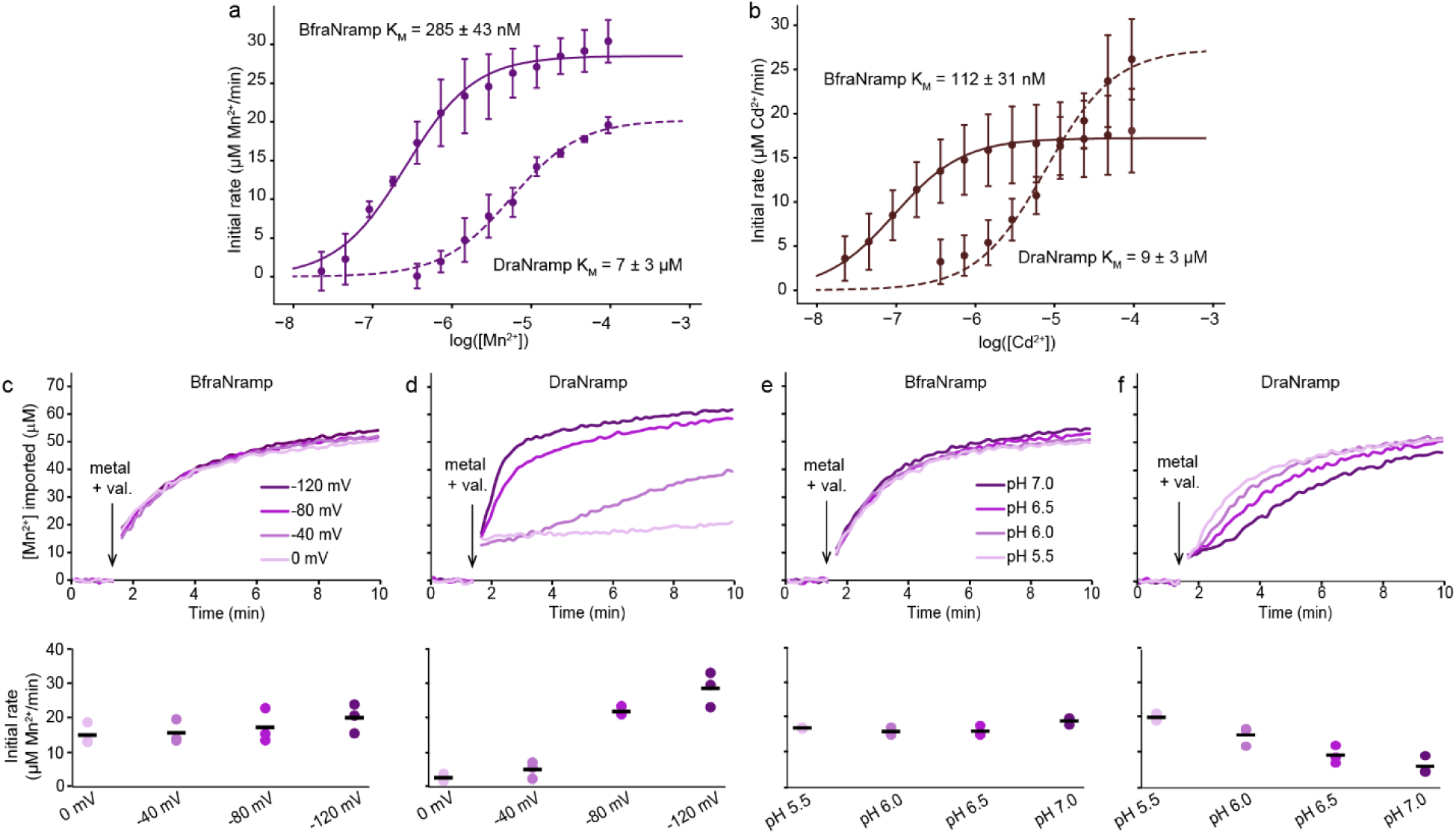
BfraNramp is a high-affinity transporter of transition metals with little voltage or proton dependence. (**a**-**b**) Dose-dependent (0.02–750 µM) transport of Mn^2+^ (**a**) and Cd^2+^ (**b**) by BfraNramp and DraNramp at a membrane potential of –120 mV and extra– and intraliposomal pH of 7.0. Solid and dashed lines represent fits for metal transport by BfraNramp and DraNramp respectively. BfraNramp transports both metals with higher apparent affinity (K_M_ = 285 ± 43 nM for Mn^2+^ and 112 ± 31 nM for Cd^2+^) than DraNramp (K_M_ = 7 ± 3 µM for Mn^2+^ and 9 ± 3 µM for Cd^2+^) as indicated by the resulting K_M_ values (Table 1). (**c**-**d**) Sample time courses (top) and corresponding initial rate dot plots (bottom) of the voltage dependence of Mn^2+^ transport, with no apparent voltage dependence for BfraNramp (**c**) and increased transport for DraNramp (**d**) as the membrane potential decreases from 0 mV to –120 mV. (**e**-**f**) Sample time courses (top) and corresponding initial rate dot plots (bottom) of the pH dependence of Mn^2+^ transport at –120mV membrane potential. BfraNramp shows no pH dependence between pH 7.0 to 5.5, and DraNramp shows increased Mn^2+^ transport as the pH becomes more acidic. For all transport experiments, Mn^2+^ (750 µM) and valinomycin were added at 90 s, as indicated. Experiments were performed in triplicate; black lines on the dot plots indicate the means. Supplementary Figure 2h shows the corresponding empty liposome data.

**Table 1.**
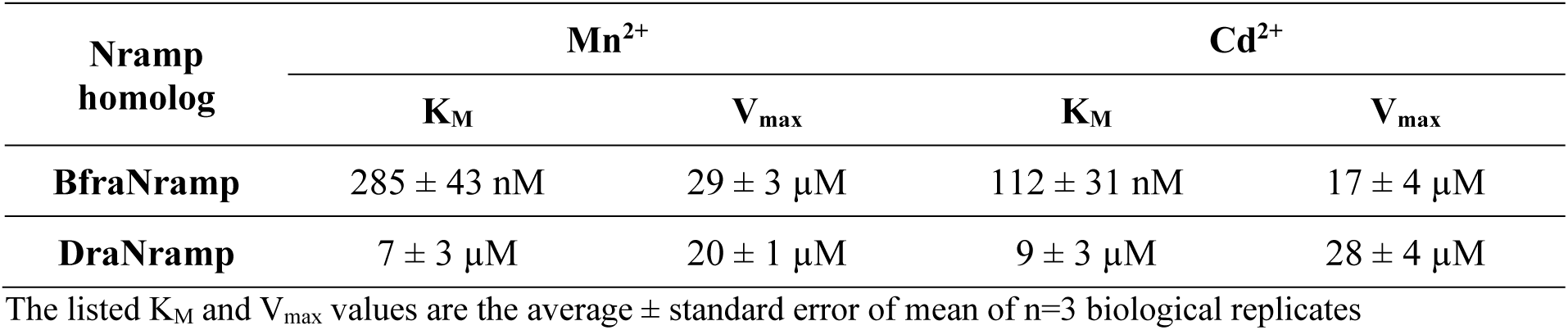
Kinetic parameters of metal transport of BfraNramp and DraNramp.

### BfraNramp does not transport protons

BfraNramp’s pH independence inspired us to test whether it can transport protons in either a metal-dependent or independent manner. Our sequence alignment reveals that BfraNramp, as a clade B Nramp, lacks most of the conserved proton-pathway residues of canonical Nramps that have been previously shown to be important for proton transport in clade A and clade C bacterial as well as eukaryotic Nramps (Fig. 1c)^5,10,14^. The pH independence of BfraNramp’s metal transport further suggests that BfraNramp may not transport protons in a metal-dependent manner. To test this hypothesis, we reconstituted BfraNramp into proteoliposomes loaded with a proton sensitive dye, BCECF^19,20^. BfraNramp did not transport protons either in the presence (Mn^2+^ or Cd^2+^) or absence of metals (Fig. 3a-b). This result contrasts with the behavior of other characterized canonical Nramps, all of which show metal-independent proton transport that is stimulated by metal ion substrate^2,5–7,10,14^, as we reproduced for DraNramp (Fig. 3a-b). The only characterized member of the broader Nramp family (Fig. 1) known to transport Mn^2+^ but not protons is EleNRMT^15^. EleNRMT is part of the bacterial ‘NRMT-like’ family that is a sister group to the canonical Nramps and whose members lack the canonical metal-binding motif as well as the conserved proton-pathway residues (Fig. 1c).

**Figure 3.**
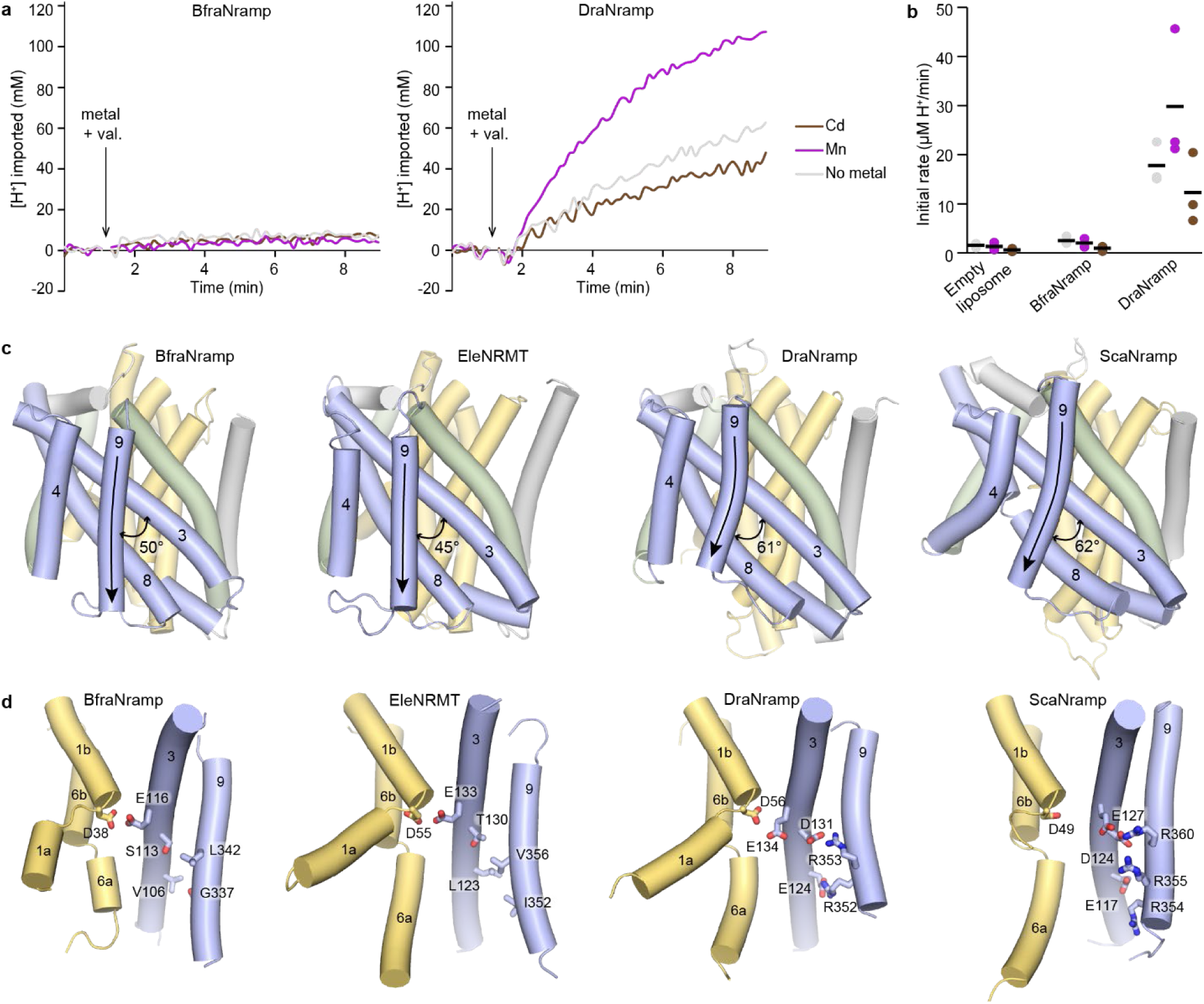
BfraNramp shows distinct proton transport behavior different from other cannonical Nramps. (**a**) Representative time courses from proteoliposome-based proton transport assay using proton-sensitive BCECF dye. BfraNramp (left) does not transport protons in the absence of metal ions or in presence of Mn^2+^ or Cd^2+^. DraNramp (right) transports protons in the absence of metal ions, and proton transport is enhanced in presence of Mn^2+^ and inhibited by Cd^2+^. (**b**) Dot plots of initial rates corresponding to proton transport for BfraNramp and DraNramp. The black lines indicate the means from triplicate measurements. **(c)** Structural comparison of different Nramps in inward-open conformation—BfraNramp (10KY), EleNRMT (7QJJ), DraNramp (8E6M), and ScaNramp (5M95) from left to right. The bundle helices (TMs 1, 2, 6, 7) are yellow, the hash (TMs 3, 4, 8, 9, labeled) blue, the arms (TMs 5, 10) green, and other structural elements are grey. The arrows highlight the curve in TM9 of the proton-transporting DraNramp and ScaNramp. (**d**) TMs 1, 2, 3, and 9 (similar to the cartoon in Fig. 1b) for the same 4 structures, highlighting in sticks the residues corresponding to the conserved proton-pathway residues in DraNramp. BraNramp and EleNRMT, both of which do not transport protons, have primarily hydrophobic residues at these positions, compared to the hydrophilic pathway and salt bridges present in DraNramp and ScaNramp, both of which transport protons.

To further investigate is distinct proton and metal transport behavior, we determined the crystal structure of BfraNramp at 2.48 Å (Table 2). In this structure, BfraNramp adopts an inward-open conformation with similar overall topology and architecture to DraNramp (Supplementary Fig. 3a; RMSD of 2.2 Å over 360 residues with the inward-open structure of WT DraNramp [PDB ID: 8E6M]). The key differences between BfraNramp and DraNramp are in the orientations of TMs 3, 4 and 9 (Fig. 3c), part of the hash that behaves as a rigid body across the transport cycle of DraNramp^12^. The most pronounced difference is in how TM9 orients itself with respect to TM3. The BfraNramp TM9 is longer, less curved, and crosses TM3 at a more acute angle than in DraNramp (50° vs. 61° overall TM3-TM9 crossing angle in BfraNramp and DraNramp, respectively; Fig. 3c). Interestingly, TM3 and TM9 harbor most of the proton-pathway residues conserved in Nramps that transport protons (Figs. 1b and 3d). TM9 of EleNRMT^15^, which like BfraNramp does not transport protons and lacks the conserved proton-pathway residues, has a similarly straighter TM9 oriented at 45° to its TM3 (Fig. 3c). In comparison, TM9 of another canonical Nramp, *Staphylococcus capitis* (Sca)Nramp from clade C^8,12^, is more similar to TM9 of DraNramp (62° angle between TM3 and TM9), with the bend accommodating the bulky arginines that form salt-bridge interactions with the TM3 acidic residues crucial for proton transport (Fig. 3d). The TM8-TM9 loop is also longer in BfraNramp and EleNRMT than in other canonical Nramps and interacts TM4 (Supplementary Fig. 3b). Overall, TMs 3, 4, 8 and 9, which contain the proton pathway in canonical Nramps and remain relatively rigid across the Mn^2+^ transport cycle, pack more closely and through more hydrophobic interactions in BfraNramp and EleNRMT. This is consistent with the hashes of BfraNramp and EleNRMT not being conducive for proton transport^13,14^. While it was hypothesized that clade B and Nramp-related proteins may co-transport protons through the same path as the metal ions^3^, our results clearly indicate that BfraNramp does not transport protons at all.

**Table 2.**
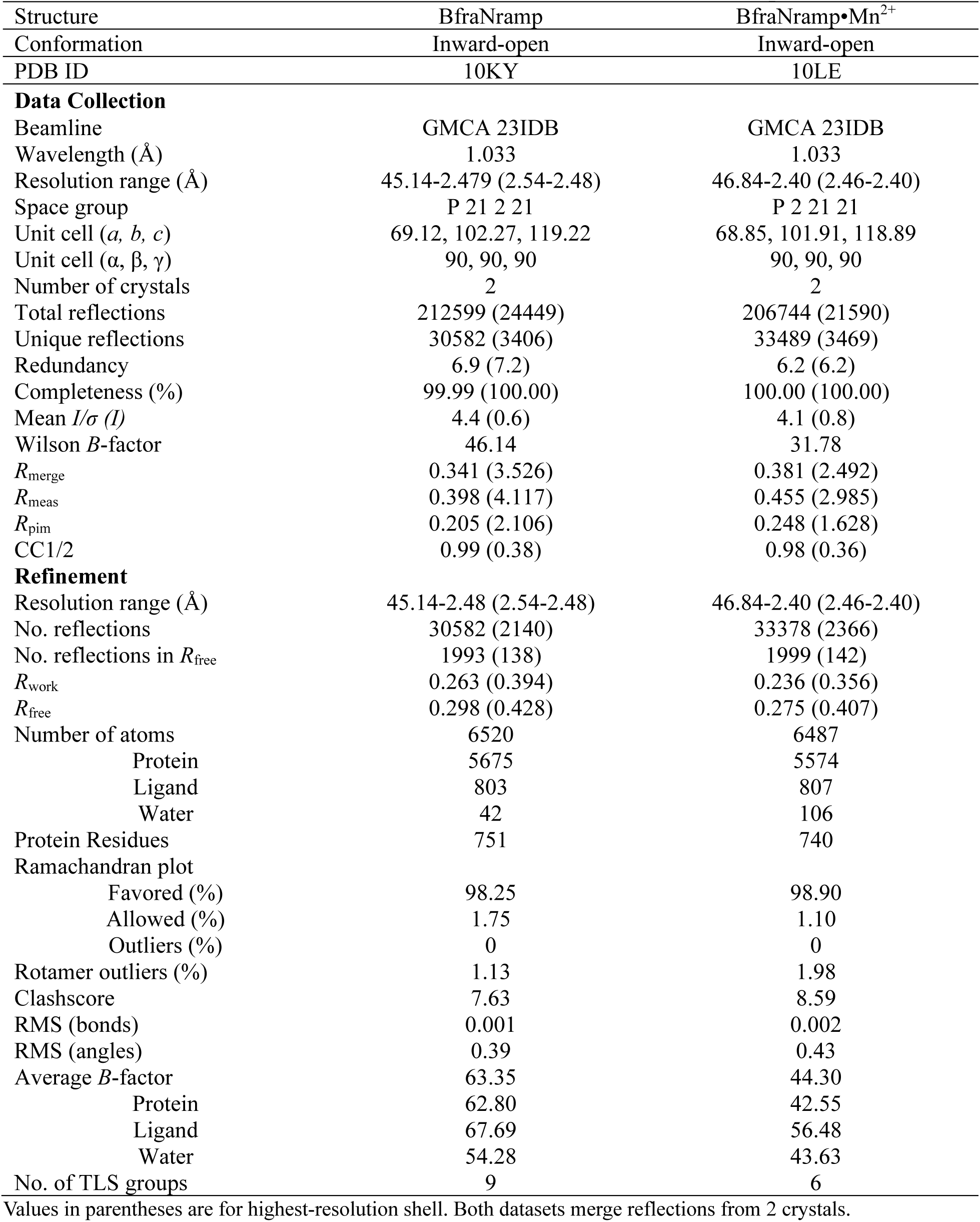
Data collection and refinement statistics for four new DraNramp structures.

We then tested whether introducing the conserved proton-pathway residues in BfraNramp would enable proton transport. Based on sequence and structure alignments with DraNramp and ScaNramp, we introduced the following mutations: S113D+L342R (mimicking the D131–R353 salt bridge in DraNramp), V106E+G337R (mimicking the E124–R352 salt bridge in DraNramp) and a quadruple mutation introducing both potential salt bridges (S113D+L342R+V106E+G337R), structurally reproducing the DraNramp proton pathway (Fig. 3d). However, none of the three variants transported protons at detectable levels in proteoliposome-based transport assays in the absence or presence of metals (Supplementary Fig. 3c-f).

These structural and biochemical results indicate that BfraNramp can robustly transport metal ions independent of any external voltage or pH driving force, and that the sequence features underlying this proton-independent transport extend beyond the previously identified proton-pathway residues. More work is required to understand how the proton co-transport behavior evolved in other canonical Nramps starting from the proton-independent clade B Nramps, as it must have been facilitated by residues outside of the known proton pathway.

### BfraNramp structure reveals a distinct metal-binding mode

BfraNramp is the only canonical Nramp family member known to not co-transport protons. Thus, we were interested in understanding whether there are any differences in how BfraNramp binds metals. We determined a 2.40-Å crystal structure of BfraNramp bound to Mn^2+^ (Table 2). BfraNramp•Mn^2+^ adopts an inward-open conformation almost identical to the metal-free structure (RSMD = 0.27 Å; Supplementary Fig. 4a). Mn^2+^ binds between the unwound regions of TM1 and TM6 like in other canonical Nramps (Fig. 4a, Supplementary Fig. 4c). However, the Mn^2+^ is displaced in comparison to previous Mn^2+^-bound Nramp structures such that, while it still interacts with the canonical TM1 aspartate (D38) and TM6 methionine (M210), it does not directly interact with the canonical TM1 asparagine (N41; distance of 6.1 Å; Fig. 4a, Supplementary Fig. 4b). Instead, Mn^2+^ directly interacts with E116 on TM3 (3.4 Å distance), a glutamate conserved in all canonical Nramps and some Nramp-related proteins (Fig. 4a, Supplementary Fig. 4c; Fig. 1c). The corresponding glutamate in DraNramp (E134) is essential for proton but not metal transport^13,14^. BfraNramp H212, the conserved histidine in the TM6 unwound region, is also close enough (3.5 Å) to directly coordinate the Mn^2+^. This histidine has been implicated in proton transport in Nramps^5^. In addition to D28, M210, E116 and H212, there are waters that complete the coordination sphere of Mn^2+^ in the inward-open BfraNramp conformation (Fig. 4a, Supplementary Fig. 4c). This difference in the Mn^2+^ position between BfraNamp and DraNramp is not unprecedented, as a different Mn^2+^ position was also observed in the cryo-EM structure of human (Hs)NRAMP2^10^. In HsNRAMP2, Mn^2+^ interacts with the canonical binding-site aspartate (D115), asparagine (N118) and methionine (M294), along with S189 and Q192 of TM3 (Fig. 4a). Superposition of the three structures shows the subtle differences in the Mn^2+^ positioning that results in distinct coordination spheres (Fig. 4a and Supplementary Fig. 4b). It remains unclear whether these differences represent distinct steps along the transport cycle or more global variations in substrate interactions in different Nramps.

**Figure 4.**
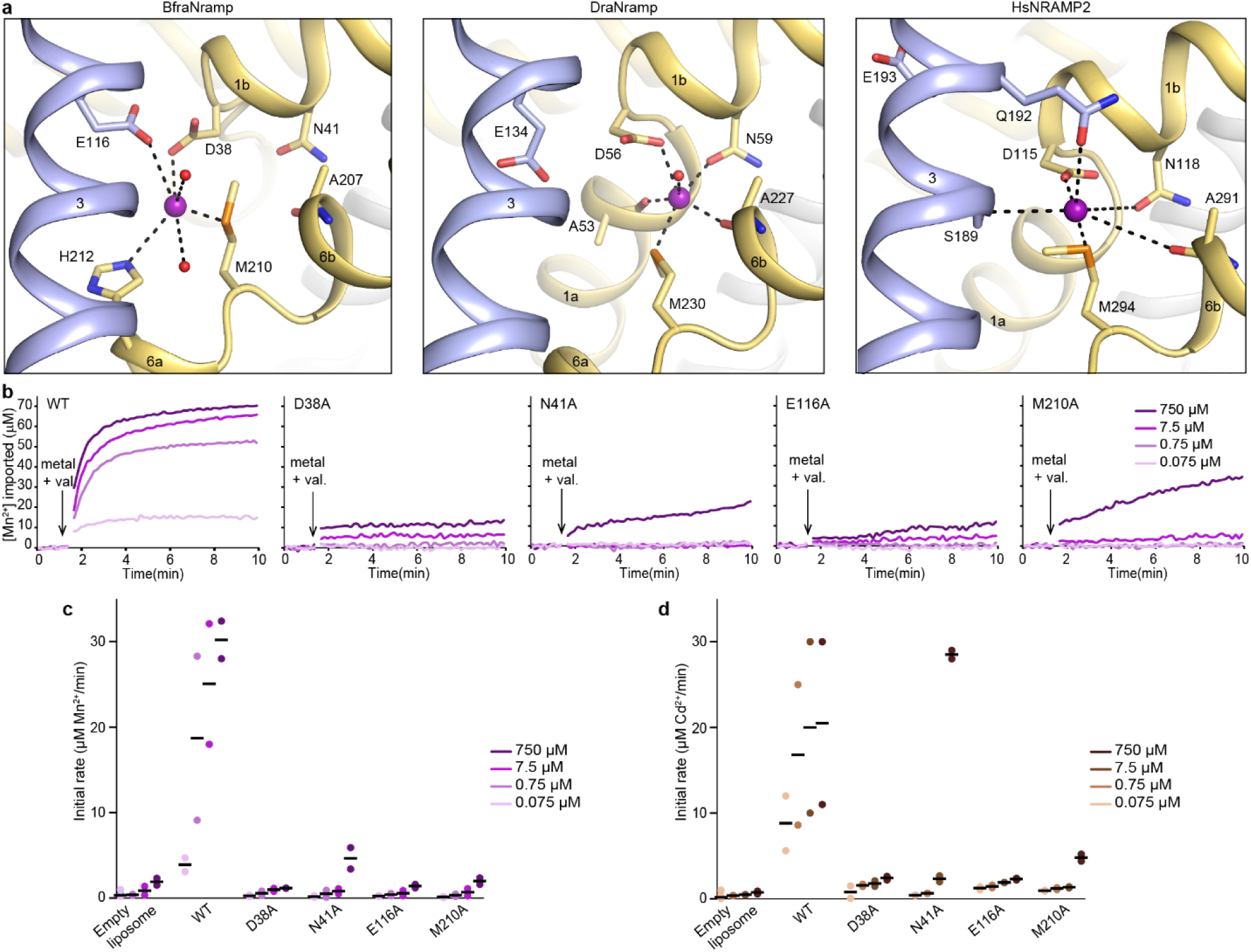
Analysis of the Mn^2+^-binding site of BfraNramp. (**a**) View of the Mn^2+^-binding site of BfraNramp (PDB ID: 10LE) in comparison to DraNramp (PDB ID: 8E60) and HsNRAMP2 (PDB ID: 9F6N). The Mn^2+^ is shifted towards TM3 and directly coordinates with E116, a residue conserved in all canonical Nramps and involved in proton transport^10,13,14^. Furthermore, in BfraNramp the Mn^2+^ ion does not bind N41, a conventional metal-coordinating residue in other canonical Nramps^8,12^. In HsNRAMP2, the Mn^2+^ is also shifted from the canonical metal-binding position^10^, although somewhat differently than in BraNramp. (**b**) Representative time courses from proteoliposome-based transport assays at different Mn^2+^ concentrations for BfraNramp variants with mutations at Mn^2+^-coordinating residues. Mutations E116A and D38A drastically impaired Mn^2+^ transport at all concentrations, whereas N41A and M210A retained some transport at the highest metal concentration (750 µM). (**c**-**d**) Dot plots of initial rates of Mn^2+^ transport (**c**; from data as in panel **b**) or Cd^2+^ transport (**d**; from data as in Supplementary Fig. 4d). Data were collected in duplicate and black bars represent the means.

We hypothesized that the glutamate conserved across the canonical Nramps clades (E116 in BfraNramp; Fig. 1c), may play distinct roles in the proton-transporting members (where it is important for proton transport) and the proton-insensitive members (directly binding Mn^2+^ in BfraNramp). Correspondingly, the E116A mutation in BfraNramp nearly eliminated transport of both Mn^2+^ and Cd^2+^ at all concentrations compared to the WT (Fig. 4b-d and Supplementary Fig. 4d). This contrasts with the analogous E134A DraNramp variant, which transports both Mn^2+^ and Cd^2+^ nearly as well as WT, although without any pH and voltage dependence unlike WT DraNramp^13,14^. Thus, the conserved glutamate plays distinct physiological roles in different Nramps.

We also tested the effect of alanine mutations at the other Mn^2+^-coordinating residues. The D38A BfraNramp variant, similar to E116A, transports essentially no Mn^2+^ or Cd^2+^ (Fig. 4b-4d and Supplementary Fig. 4d). Both M210A and N41A only showed transport activity at the highest concentration (750 µM) of both metals, with only N41A reaching near-WT levels of Cd^2+^ transport at that concentration, suggesting a much poorer affinity for these metal ion substrates (Fig. 4b-4d and Supplementary Fig. 4d). The deleterious effect of N41A on Mn^2+^ transport contrasts with its lack of a direct interaction with Mn^2+^ in the BfraNramp•Mn^2+^ structure. These results indicate that N41 is important for transporter function, either indirectly stabilizing the bound Mn^2+^ substrate or interacting directly with it in another conformation along the transport cycle.

## Discussion

Our biochemical and structural analyses of BfraNramp, a canonical Nramp from the basal bacterial clade B, reveal a distinctive proton-independent metal transport mechanism. Unlike other characterized canonical Nramps^5,8,10,14^, BfraNramp transports its likely physiological substrate, Mn^2+^, without any external energy drive from the membrane potential or protons. BfraNramp also does not transport protons in the absence of metals, unlike other canonical Nramps which have metal-independent proton leaks^2,6,7,9,13,14^. Our evolutionary analysis highlights that this correlates with the clade B Nramps lacking most of the conserved proton-pathway residues found in other clades of canonical Nramps^13,14^. In our structure of Mn^2+^-bound BfraNramp in an inward-open state the Mn^2+^ position is shifted compared to most Nramps, enabling a direct interaction with E116, a glutamate conserved across canonical Nramps.

Our structure-guided evolutionary analysis agrees with previous phylogenetic classification of the Nramp superfamily^3,4^ and positions clade B Nramps, including BfraNramp, at the base of the canonical bacterial Nramp branch from the paraphyletic Nramp-related proteins. The Nramp-related proteins have a diverse amino acid composition in place of the metal-coordinating DPGN (TM1) and MPH (TM6) motifs conserved in canonical Nramps. They also lack the proton-pathway residues in TMs 3 and 9 conserved in the proton-transporting canonical Nramps (i.e., excluding bacterial clade B); these are instead replaced by largely non-polar residues. Clade B thus likely serves as a living remnant of an evolutionary intermediate, possessing the conserved metal-binding DPGN and MPH motifs but lacking the charged proton-pathway residues.

It has been assumed that the ancestral clade B Nramps would co-transport protons similarly to the other, derived, canonical Nramps^3,4^. Our finding that BfraNramp does not transport protons instead supports the idea that clade B Nramps are transition metal ion uniporters, more mechanistically similar to the Nramp-related protein EleNRMT, a Mg^2+^ transporter that does not co-transport protons^15^. BfraNramp has better apparent affinity for Mn^2+^ than DraNramp (K_M_ ∼ 300 nM and 7 µM, respectively). The published K_M_ values of other canonical Nramps are most similar to that of DraNramp (EcoDMT, 18 µM; HsNRAMP1, 4 µM; and HsNRAMP2, 7 µM)^5,12,13^. BfraNramp’s robust Mn^2+^ transport activity excludes the possibility that its proton co-transport activity is simply too low to be measured, as we can readily detect proton co-transport by DraNramp in the same assay. Its faster kinetics at low metal concentrations suggest that BfraNramp has lower energy barriers for the outward-inward conformational switches. This is reminiscent of a recent finding that prokaryotic glutamate transporters evolved from proton-driven to sodium-driven transporters through allosteric mutations, yielding an uncoupled transporter intermediate^21^. Our combined functional and phylogenetic analyses indicate that clade B represents an evolutionary intermediate in the Nramp phylogeny—characterized by a transition metal-binding motif without ion coupling—and that this step was then followed by the emergence of proton-dependent metal transport, giving rise to the other canonical Nramp clades.

The Mn^2+^-bound structure of BfraNramp adds a new layer to our understanding of metal binding and transport by Nramps. Mn^2+^ binds between TMs 1 and 6, as in other canonical Nramps, but it is shifted towards TM3 where it interacts with E116 rather than N41 on TM1. This observation aligns with our mutagenesis results: E116A abolished metal transport similarly to the D38A mutation to the canonical TM1 aspartate, whereas N41A was less deleterious (and less so than the analogous N59A mutation in DraNramp^16^). The Mn^2+^ ion is also displaced in the HsNRAMP2 structure, although it contacts TM10 rather than TM3^10^. These variations in metal coordination suggest subtle mechanistic differences superimposed on an overall similar global transport cycle for canonical Nramps. E116 is conserved across canonical Nramps and appears as glutamate, aspartate, or asparagine in several Nramp-related protein clades (Fig. 1c). In proton-coupled Nramps, this residue is crucial for proton transport^5,13^. Our results suggest that it originally evolved to support metal transport, and only later gained a role in proton co-transport as the location of the metal-binding site shifted away from it and it took on a new function.

Overall, our work fleshes out the evolutionary history of Nramps by functionally and structurally characterizing a member of the bacterial clade B. The Nramp-related proteins are a paraphyletic group that includes alkaline earth metal ion transporters lacking coupled driving ions, like EleNRMT. The clade B Nramps emerged as the first canonical transition metal transporters in the superfamily, likewise operating without ion coupling. Proton co-transport subsequently emerged in clade A Nramps, with a polar proton pathway separate from that of the transported metal ion. The existence of highly efficient transporters like BfraNramp that function without proton co-transport underscores that the selective advantage of proton co-transport in bacterial clades A and C and eukaryotic Nramps is not yet fully explained. Given that Nramps transport transition metals down favorable electrochemical gradients, proton co-transport likely confers benefits beyond basic transport efficiency. Well-characterized homologs from each clade now provide a foundation for comparative complementation studies linking the evolution of proton coupling and voltage dependence to physiological adaptation.

## Methods

### Multiple sequence alignment

We structurally aligned eight Nramp and NSS proteins based on six high-resolution experimental structures and two AlphaFold models (Supplementary Table 1). We used this structural alignment to generate a structure-based sequence alignment with Promals3D^22^ but could identify by eye a large number of misalignments from the superposed structures, and so we manually modified this alignment to better match the aligned structures. To assemble a larger set of sequences, we ran Jackhmmer on the UniprotKB database for five iterations with a bitscore threshold of 50 starting from five separate sequences—DraNramp (UniProt ID: Q9RTP8), EleNRMT (UniProt ID: C8WJ67), a second Nramp-related protein from *Eggerthella lenta* (UniParc ID: UPI0002171069), *Acinetobacter baumannii* MumT (A0A385EU17), and *Aquifex aeolicus* LeuT (UniProt ID: O67854)—and merged the complete sequences collected from these runs, yielding a total of 109,451 sequences. We then used HMMER tools^23^ to build an HMM profile from the eight-sequence structural alignment and used that HMM profile to align the 109,451 sequences. Most sequences aligned well to this profile, with relatively few gaps in the core TMs, while 10,512 sequences were removed from the final alignment for not aligning well to the core helical domain, leaving 97,939 sequences for downstream analyses.

### Phylogenetic inference

To understand the phylogenetic relationships between these sequences, we used a subset of 69 highly diverse sequences of interest to infer the posterior probabilities of various tree topologies with MrBayes^24^ v3.2.7. We included the 435 alignment columns whose average alignment probability score, as determined by HMMalign, was greater than 50%. MrBayes was run with 4 chains for 1,000,000 iterations using inverse-gamma distributed rate parameters, a mixed amino acid model prior, and all other default priors. The python library ete3^25^ was used to analyze posterior probabilities of various sub-topologies of the tree, and a majority-rule consensus tree was generated with MrBayes and visualized in Geneious. We used the maximum a posteriori tree in MrBayes (shown in Fig. 1a) to constrain the topology of a larger approximately maximum-likelihood build using FastTree^26^ v2.1.11 with the complete alignment filtered down to only Uniref90 sequences, totaling 36,700. Clades in the tree were identified manually and sequence motifs were generated using logomaker^27^ in Python.

### Cloning of BfraNramp constructs

The WT BfraNramp sequence, except with an N-terminal truncation of 13 residues (ΔN13), was cloned into a modified pET21a vector with an N-terminal 8xHis-tag^16^ using the following primers: 5’– GCACATATGAAACGCTATCTGGGAGGTTTGGAC –3’ and 5’-ATGTGCGGCCGCTTACGAGAACAACAACATTATATTCAACACTGT –3’. All mutations were made on this ΔN13 background, which we refer to as WT throughout this paper, in the same vector using a Quikchange mutagenesis protocol (Stratagene) with the primers listed in Supplementary Table 2 and confirmed by Sanger DNA sequencing.

### Protein expression

WT and mutated BfraNramp proteins were expressed in *E. coli* C41(DE3). Saturated overnight cultures were used to inoculate at 1:100 dilution terrific broth supplemented with 10% (wt/vol) glycerol and 100 mg/L ampicillin. Cells were grown at 37°C to OD_600_ = 0.9, and then the temperature was reduced to 18°C and cells were induced with 0.5 mM IPTG and grown for 16 h, harvested, either proceeding with purification, or flash frozen in liquid nitrogen and stored at −80°C for later use.

### Protein purification for crystallography

For crystallography, WT BfraNramp-expressing cells harvested from 10 L of culture were resuspended in load buffer (20 mM sodium phosphate, pH 7.5, 55 mM imidazole pH 7.5, 500 mM NaCl, 10% (v/v) glycerol) supplemented with 1 mM PMSF, 1 mM benzamidine, 0.3 mg/mL DNAse I and 0.3 mg/mL lysozyme in 2 mL of load buffer per g of cells, incubated 30 min 4°C, then lysed by sonication on ice (5-6 cycles of 45 s with a Branson Sonifier 450 under duty cycle of 65% and output 10). Lysates were cleared of cell debris by centrifuging for 20 min at 27,200 × *g* in a JA-20 rotor (Beckman) and membranes were pelleted from the clarified lysate by ultracentrifugation at 235,000 × *g* in a Ti-45 rotor (Beckman Coulter) for 70 min. Membranes from 70 mL of clarified lysate were homogenized in 40 mL load buffer using a glass Potter-Elvehjem grinder, solubilized for 1 h in 1% (w/v) n-dodecyl-β-D-maltopyranoside (DDM), then ultracentrifuged at 142,000 × *g* in a Ti-45 rotor for 35 min to remove insoluble materials. The supernatant was loaded onto Ni-Sepharose resin (GE Healthcare) pre-equilibrated with load buffer (0.5 mL resin per liter of cells) and nutated for 90 min at 4°C in, then washed in a batch format with 20 column volumes (CV) of each of the following three buffers sequentially: (i) load buffer containing 0.03% DDM, (ii) load buffer containing 0.5% lauryl maltose neopentyl glycol (LMNG), and (iii) load buffer containing 0.1% LMNG. Protein was eluted in 10 CV of 20 mM sodium phosphate, pH 7.5, 450 mM imidazole pH 7.5, 500 mM NaCl, 10% (v/v) glycerol, 0.01% LMNG. Protein elution was confirmed by Coomassie-stained SDS-PAGE, and then the eluate concentrated to < 0.5 mL in a 50 kDa molecular weight cutoff (MWCO) centrifugal concentrator (EMD Millipore) and purified on a Superdex S200 10/300 (GE Healthcare) pre-equilibrated (10 mM HEPES pH 7.5, 150 mM NaCl, 0.003% LMNG). Peak protein fractions were confirmed by Coomassie-stained SDS-PAGE, pooled, concentrated to ∼25 mg/mL using a 50 kDa MWCO centrifugal concentrator, aliquoted, and flash frozen in liquid nitrogen and stored at –80°C. Protein from several purifications (n=3) resulted in similar crystals.

### Purification of BfraNramp for in vitro transport assays

To purify WT and mutant BfraNramp for proteoliposome-based transport assays, harvested cells from 6 L of culture were lysed, sonicated and membranes were isolated, homogenized and solubilized in 1% DDM and loaded to Ni-Sepharose resin (GE Healthcare) using the same protocol as for crystallization. The resin was washed thrice in a batch format with 20 CV of load buffer with 0.03% DDM. Bound protein was eluted in 10 CV of 20 mM sodium phosphate, pH 7.5, 450 mM imidazole pH 7.5, 500 mM NaCl, 10% (v/v) glycerol, 0.03% DDM. The eluted protein was confirmed by SDS-PAGE and concentrated to ∼2.5 mL using a 50 kDa MWCO centrifugal concentrator, then buffer-exchanged into 150 mM NaCl, 10 mM HEPES, pH 7.5, and 0.1% (wt/vol) β-decylmaltoside (DM) using PD-10 desalting columns (GE healthcare). Protein was concentrated to ∼1 mg/mL using a 50 kDa MWCO centrifugal concentrator and flash frozen in liquid nitrogen and stored at –80°C. Purifications of each protein construct were performed at least twice and resulted in similar data.

### Proteoliposome-based in vitro transport assays

Liposome preparation and metal transport assays were performed using similar protocols as previously described^12–14,16,28^. Briefly, lipids (3:1 mass ratio of 1-Palmitoyl-2-oleoyl-sn-glycero-3-phosphoethanolamine (POPE): 1-Palmitoyl-2-oleoyl-sn-glycero-3-phospho-rac-glycerol (POPG); Avanti Polar Lipids) were dried, resuspended in buffer containing 90 mM KCl, 30 mM NaCl, 10 mM MOPS pH 7.0 and 5 mM DM, and homogenized in a bath sonicator. BfraNramp was mixed with the prepared lipids at a 1:400 mass ratio and the mixture dialyzed against 90 mM KCl, 30 mM NaCl, 10 mM MOPS pH 7.0, 0.2 mM EDTA at 4°C for 1 day, 90 mM KCl, 30 mM NaCl, 10 mM MOPS pH 7.0, 0.1 mM EDTA at 4°C for 1 day, then 90 mM KCl, 30 mM NaCl, 10 mM MOPS pH 7.0 at room temperature for 1 day. EDTA was used in the initial buffers to remove any residual metals. Metal-sensitive dye Fura-2 (100 μM) or proton-sensitive dye BCECF (150 μM) was incorporated into the proteoliposomes post dialysis using three freeze-thaw cycles. The dye-loaded proteoliposomes were flash-frozen and stored at –80°C for further use.

On the day of the assay, the liposomes were extruded through a 400-nm filter, separated from bulk dye on a PD-10 column (GE Life Sciences) and diluted fivefold in buffer containing 0.5 mM MOPS pH 7.0, and 100 mM NaCl in 96-well black clear-bottom assay plate (Greiner). For experiments at different voltages, buffers were prepared with varying amounts KCl and NaCl to create different membrane potentials in the presence of valinomycin^28^. For experiments at lower extraliposomal pH values, buffers were prepared containing 0.5 mM MES (pH 5.5-6.5) at respective voltages^28^. Metal transport was monitored by measuring Fura-2 fluorescence at λ_ex_ = 340 and 380 nm, at λ_em_ = 510 nm and proton transport was monitored by measuring BCECF fluorescence at λ_ex_ = 450 and 490 nm, at λ_em_ = 535 nm. Both metal and proton transport assays were performed at room temperature. Baseline fluorescence was measured for 90 s, then metal substrate was added along with 50 nM valinomycin to establish a negative internal potential and fluorescence further monitored for 600 s. The transport data were analyzed and plotted in Excel as described^28^. For each construct, the presented data originates from 2-3 batches of reconstituted liposomes, each from an independent protein purification, represented as scatterplots and the corresponding means.

### BfraNramp crystallization

Metal-free and Mn^2+^-bound BfraNramp were crystallized in lipidic cubic phase (LCP) where protein was mixed with monoolein in 1:1.5 volume ratio using the syringe reconstitution method^13^. The protein bolus and precipitant (0.2 M ammonium sulfate, 0.02 M sodium chloride, 0.02 M sodium acetate pH 4.0, 33% PEG 200) were dispensed in a 1:12 v/v ratio onto custom-made 96 well glass sandwich plates using an NT8 drop-setting robot (Formulatrix). Crystals (∼50-μm rods) were harvested within 5-7 days on MicroMesh mounts (MiTeGen) and flash-frozen in liquid nitrogen prior to data collection.

### ***X-*** ray diffraction data collection and processing

Beamline 23-ID-B of the Advanced Photon Source at wavelength of 1.033 Å was used for diffraction data collection. The crystals were located in the meshes by grid scanning with a 20-μm beam at 10% transmission followed by data collection with a 10-μm beam at 15% transmission. Data were indexed done in XDS^29^ and scaled in CCP4 AIMLESS^30,31^ version 7.0. For both metal-free and Mn^2+^-bound structures, datasets collected from two crystals were independently indexed and integrated, then combined during scaling using CCP4 AIMLESS^30,31^ to obtain complete datasets. The resolution cutoff of each dataset was based on CC_1/2_ values of 0.3 and above^32,33^. Initial phases were determined by molecular replacement in PHENIX^34^ version 1.17.1-3660 using a BfraNramp model generated by Alphafold2^35^ as search model. Data statistics are listed in Table 2.

### Model building, refinement, and analysis

Models were built in COOT^36^ version 7.0 and refined in PHENIX^34^, with refinement cycles including reciprocal space, TLS groups, and individual B-factor refinement, and optimization of the X-ray/stereochemistry and X-ray/ADP weights. Ligand restraints for monoolein and PEG were generated in Phenix.elbow with automatic geometry optimization. Structures contain two protein molecules in the asymmetric unit. The metal-free structure spans residues 29-417, except residues 221–233 in chain A and 221–234 in chain B, which lacked interpretable electron density. The Mn^2+^-bound structure spans residues 29-221, 233-325 and 334-417 in chain A and 30-221, 233-326 and 334-417 in chain B. Chains A and B superimpose with an RMSD of 0.324 Å (373 C_α_ atoms), and chain A was used for all analyses. Refinement statistics are listed in Table 2. Structural biology applications used in this project were compiled and configured by SBGrid^37^.

## Acknowledgements

We thank previous and current members of the Gaudet lab for discussions and assistance, particularly José Velilla for help with crystal soaking and fishing prior to data collection. This work was funded in part by NIH grant R01GM120996 (R.G.), the NSF-Simons Center for Mathematical and Statistical Analysis of Biology at Harvard (award number 1764269) and the Harvard Quantitative Biology Initiative (S.P.B.). Diffraction data reported in this study were collected at GM/CA beamline 23IDB in the Advanced Photon Source. GM/CA is funded by the National Cancer Institute (ACB-12002) and the National Institute of General Medical Sciences (AGM-12006, P30GM138396). The Eiger 16M detector at GM/CA-XSD is funded by NIH grant S10 OD012289. The Advanced Photon Source is a U.S. Department of Energy Facility operated by Argonne National Laboratory under Contract No. DE-AC02-06CH11357.

## Author contributions

RG and SR conceptualized the study. SR performed cloning, protein purification, crystallography, and functional assays. SB performed the structure-guided evolutionary and sequence analyses. SR, SB and RG wrote the manuscript, and all authors edited the manuscript.

## Materials & Correspondence

The correspondence and material requests should be addressed to.

