## Supplemental Tables and Figures for "Structural and functional basis of proton-independent transition metal import by a canonical bacterial Nramp transporter"

This file contains:


- Supplementary Table 1. Structures and AlphaFold models used as seeds for bioinformatics analyses
- Supplementary Table 2. Primers for BfraNramp cloning
- Supplementary Figure 1. Additional phylogenetic analysis of Nramp and Nramp-related protein sequences
- Supplementary Figure 2. Purification and transport of properties of BfraNramp
- Supplementary Figure 3. Structural and functional analysis of BfraNramp
- Supplementary Figure 4. Structural and functional analysis of BfraNramp bound to $Mn^{2+}$

**Supplementary Table 1. Structures and AlphaFold models used as seeds for bioinformatics analyses**

| Sequence name | Organism | Structure (PDB ID or AlphaFold) | UniProt ID | Family |
|---|---|---|---|---|
| **DraNramp** | *Deinococcus radiodurans* | 6BU5 | MNTH_DEIRA | Canonical Nramps |
| **EleNRMT** | *Eggerthella lenta* | 7QIA | E4KPW4_9LACT | Nramp-related magnesium transporters |
| ***E. lenta* Nramp-related transporter 2** | *Eggerthella lenta* | AlphaFold model | A0A36N1S1_EGGLN | Unknown Nramp-related transporters |
| **Uncharacterized protein** | *Haloferax mediterranei* | AlphaFold model | I3R5I0_HALMT | Unknown Nramp-related transporters |
| **Uncharacterized protein PM0681** | *Pasteurella multocida* | AlphaFold model | Y681_PASMU | Unknown Nramp-related transporters |
| **LeuT** | *Aquifex aeolicus* | 2A65 | O6754_AQUAE | Neurotransmitter-sodium symporters |
| **Serotonin transporter** | *Homo sapiens* | 5I6X | SC6A4_HUMAN | Neurotransmitter-sodium symporters |
| **Dopamine transporter** | *Drosophila melanogaster* | 4XP4 | DAT_DROME | Neurotransmitter-sodium symporters |

**Supplementary Table 2. Primers for BfraNramp cloning**

| Primer name | Forward primer (5'-3') | Reverse primer (5'-3') |
|---|---|---|
| D38A | TTTATTGCTCCGGGCAATTGGGC TTCTAAT | GCCCGGAGCAATAAAACCTACAGT AACCAA |
| N41A | CCGGGCGCTTGGGCTTCTAATTT TGCG | AGCCCAAGCGCCCGGATCAATAAA ACCTAC |
| V106E | ACAGCCGAACTTGCTTCCATCTC TACATCA | AGCAAGTTCGGCTGTCCCCAGTATG GG |
| S113D | TCTACAGATCTGGCCGAGATTCT GGGAGG | GGCCAGATCTGTAGAGATGGAAGC AAGTAC |
| E116A | CTGGCCGCGATTCTGGGAGGGGC CATA | CAGAATCGCGGCCAGTGATGTAGA GATGGA |
| M210A | GTGGTGGCGCCTCACAATCTTTT CCTGCATTCG | GTGAGGCGCCACCACAGCACCCAG CACACT |
| G337R | CAGGTAAGGGTTATCCTGTCGTT GGGCATT | GATAACCCTTACCTGAGAGTGACTA TCCTT |
| L342R | CTGTCGAGGGGCATTGCATTGCT GCTGATT | AATGCCCCTCGACAGGATAACCCCT ACCTG |

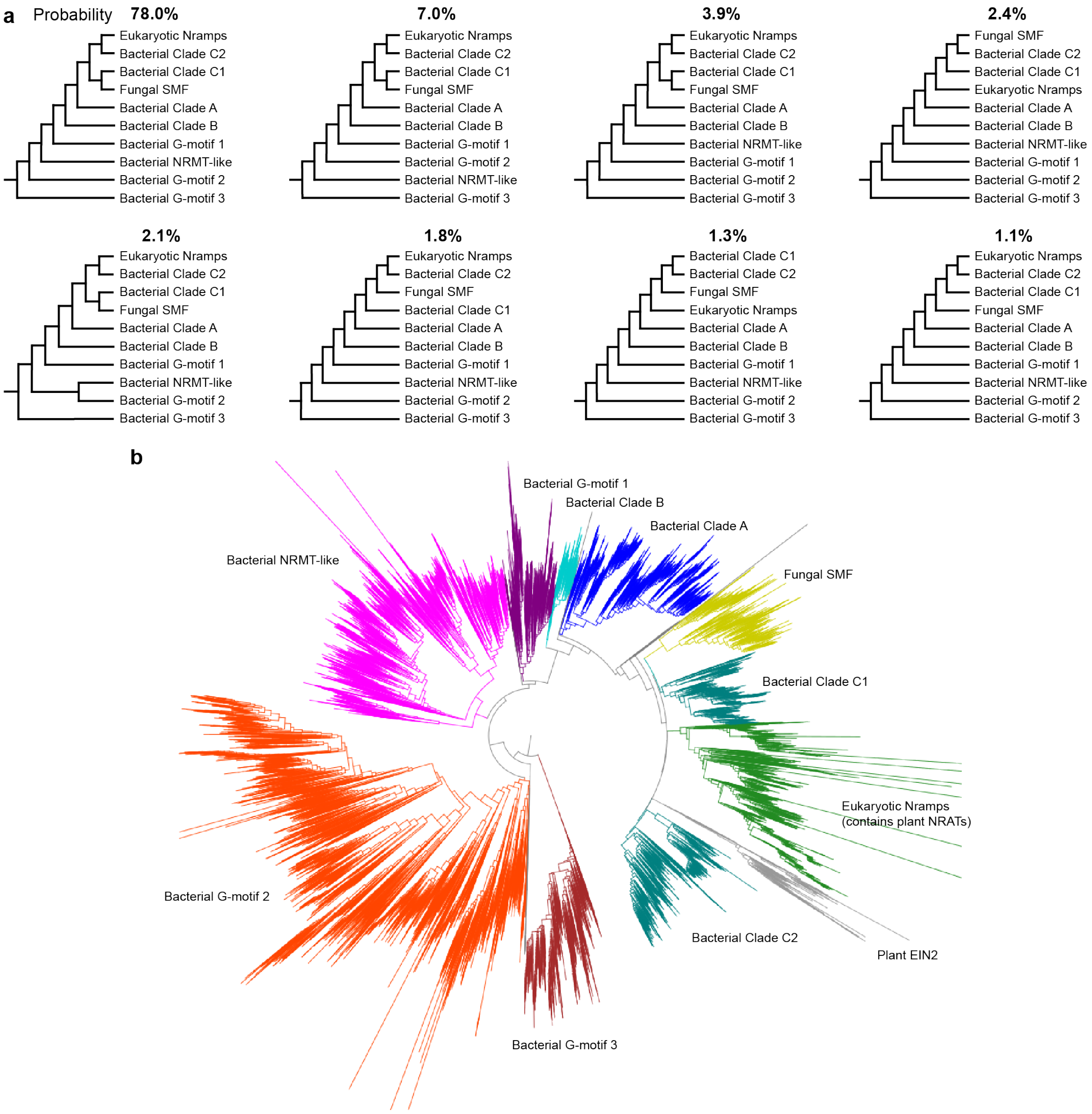


**Supplementary Figure 1. Additional phylogenetic analysis of Nramp and Nramp-related protein sequences.** (**a**) Different possible phylogenetic relationships between key sequences in the Nramp tree, with posterior probabilities of each topology estimated via MrBayes and arrangements of the ten clades with at least 1% posterior probability highlighted. All architectures share many commonalities, such as placing the bacterial G-motif clade 3 at the base of the tree. (**b**) Schematic of the entire 36,700-sequence tree inferred with FastTree constrained to the most probable MrBayes architecture, with clades highlighted as in Figure 1a.

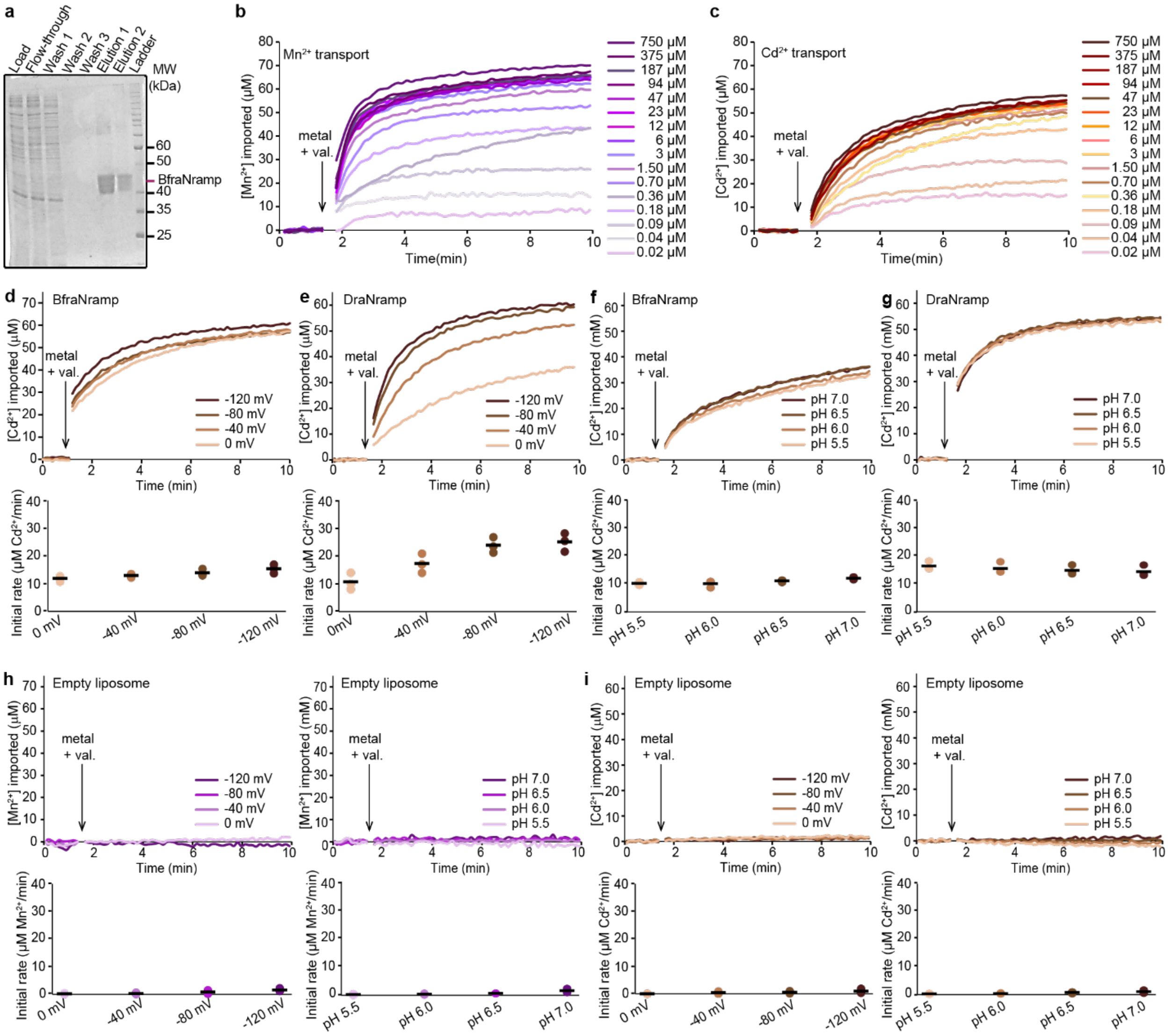


**Supplementary Figure 2. Purification and transport of properties of BfraNramp.** (**a**) Coomassie-stained 12% SDS-PAGE gel of fractions from the Ni-affinity purification of His-tagged BfraNramp. (**b-c**) Representative time courses of transport of $Mn^{2+}$ (**b**) and $Cd^{2+}$ (**c**) at different metal concentrations and -120 mV by BfraNramp in proteoliposomes. (**d-e**) Representative time courses of $Cd^{2+}$ transport (top) and corresponding initial-rate dot plots (bottom) by BfraNramp (**d**) and DraNramp (**e**) reconstituted into proteoliposomes at different membrane potentials. BfraNramp showed little voltage dependence, whereas $Cd^{2+}$ transport by DraNramp is voltage dependent. (**f-g**) Representative time courses of $Cd^{2+}$ transport (top) and corresponding initial rates (bottom) by BfraNramp (**f**) and DraNramp (**g**) at -120 mV and at different external pHs. Both BfraNramp and DraNramp show negligible pH dependence. (**h-i**) Representative time courses (top) and corresponding initial-rate dot plots (bottom) for empty liposomes for both $Mn^{2+}$ (**h**; related to Fig. 2c-f) and $Cd^{2+}$ (**i**; related to panels **d-g**). All proteoliposome experiments were performed in triplicate and the individual values represented in the dot plots, with the means indicated by black bars.

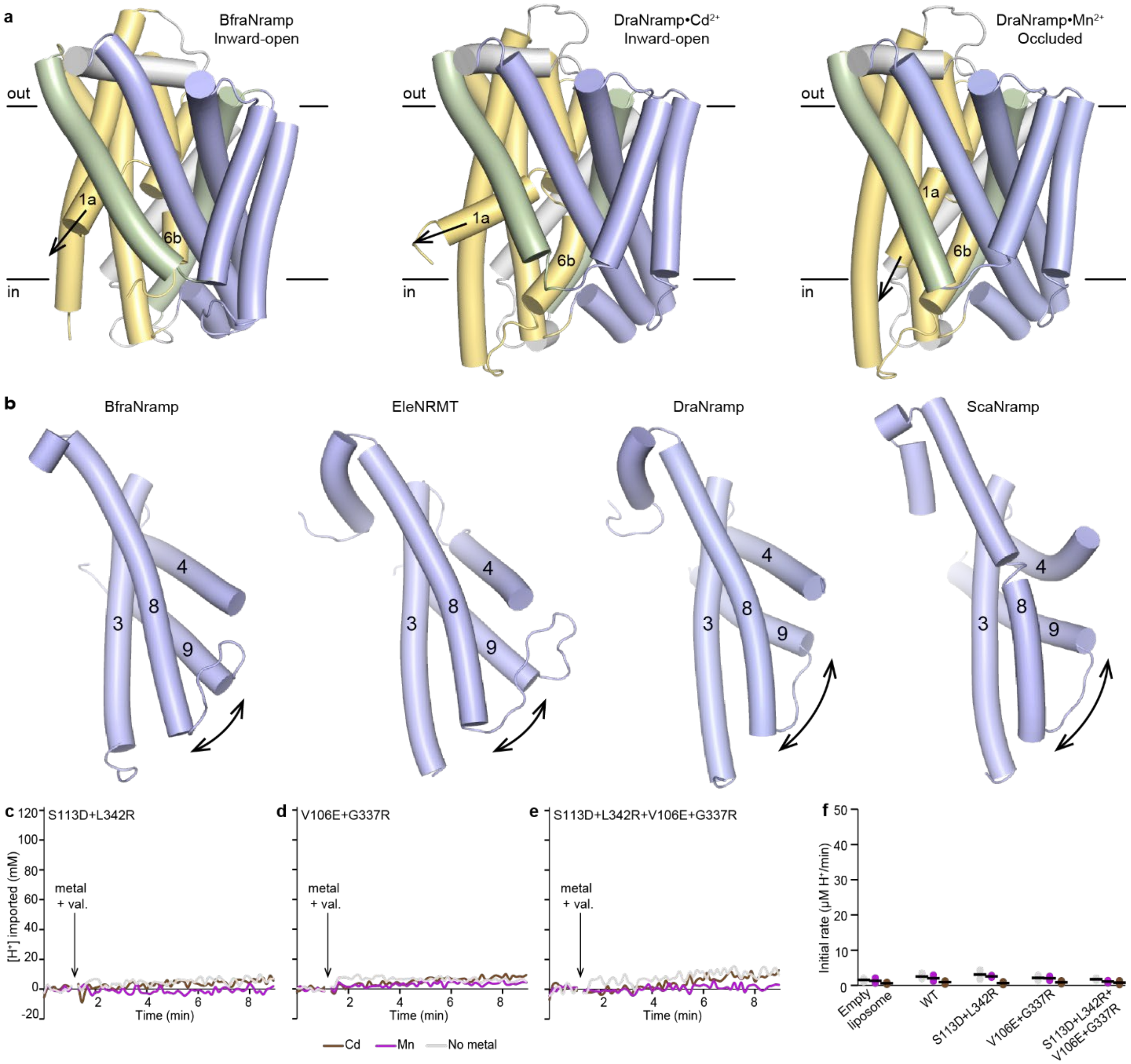


**Supplementary Figure 3. Structural and functional analysis of BfraNramp.** (**a**) Structures of BfraNramp(PDB ID: 10KY), and inward-open (PDB ID: 8E6M) and occluded (PDB ID: 8E60) conformations of DraNramp highlighting that BfraNramp is inward open. The bundle (TMs 1, 2, 6, 7) is yellow, the hash (TMs 3, 4, 8, 9) blue, the arms (TMs 5, 10) green, and other structural elements are grey. (**b**) Comparison of the hash in different inward-open Nramps. TM9 is straighter in BfraNramp and EleNRMT, and the TM8-TM9 loop is longer and different in shape in BfraNramp and EleNRMT than in the proton-transporting DraNramp and ScaNramp. (**c-f**) Representative time courses of proton transport by BfraNramp variants incorporated into proteoliposomes. WT did not transport protons in the presence or absence of metal ions (Fig. 3a). Mutations incorporating salt-bridge mimics that are crucial for proton transport in DraNramp—S113D+L342R (**c**), V180E+G337R (**d**), and S113D+L342R+V106E+G337R (**e**) did not lead to increases in proton import. **(f)** Initial proton transport rates (n = 2, bars represent mean values).

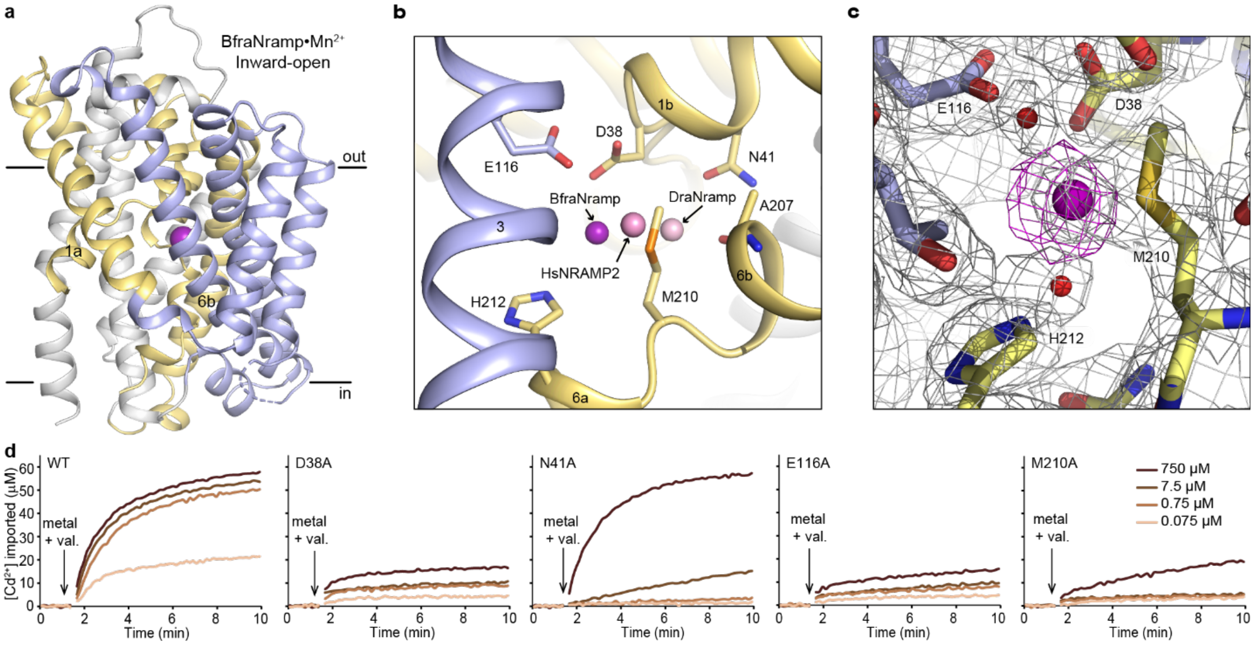


**Supplementary Figure 4. Structural and functional analysis of the BfraNramp $Mn^{2+}$ binding site.** (**a**) Structure of BfraNramp showing that $Mn^{2+}$ binds in an inward-open conformation. (**b**) BfraNramp structure showing the $Mn^{2+}$ positions in BfraNramp, DraNramp, and HsNRAMP2. (**c**) $2F_o$-$F_c$ map (gray mesh; 1σ) of the BfraNramp $Mn^{2+}$-binding site, with the omit $F_o$-$F_c$ map (magenta mesh; 2σ) corresponding to the bound $Mn^{2+}$. **(d)** Representative time courses of concentration-dependent transport of $Cd^{2+}$ by different BfraNramp variants with metal binding-site mutations. Transport is drastically impaired in D38A and E116A and partially retained in M210A and N41A, particularly at 750 μM $Cd^{2+}$. The corresponding dot plot is Figure 4d. All proteoliposome experiments were performed in duplicate and the individual values represented in the dot plots.
